# Assessing the re-introduction of *Aedes aegypti* in Europe. Will the Yellow Fever Mosquito colonize the Old World?

**DOI:** 10.1101/2020.06.04.133785

**Authors:** Daniele Da Re, Diego Montecino-Latorre, Sophie O. Vanwambeke, Matteo Marcantonio

## Abstract

*Aedes aegypti* are feared invasive mosquitoes as they transmit pathogens which cause debilitating diseases in humans. Although mainland Europe has not yet witnessed re-establishment and diffusion of *Ae. aegypti* populations, several urban areas along coastlines represent suitable habitats for the species. In addition, European coastal areas are characterized by a high exotic species propagule pressure, due to the dense international ship traffic.

Here, we applied a process-based population dynamical model to simulate both the life cycle and dispersal of *Ae. aegypti* at the local scale after its introduction through ship traffic. We selected five European ports along a gradient of latitude by considering both environmental conditions and the economical importance of ports: Algeciras and Barcelona in Spain; Venice and Genoa in Italy and Rotterdam in the Netherlands. The model was informed using parameters relevant for *Ae. aegypti* biology, fine-scale temperature time-series, urban structures and road networks.

According to model results, the introduction of small quantities of *Ae. aegypti* eggs, from 10 to 1000, has the potential to cause species establishment, high local densities and slow initial dispersal in the two southernmost study areas, Algeciras and Barcelona, whereas Genoa may be considered only close to suitability. Barcelona had the highest simulated mosquito densities (584 females/ha), whereas Algeciras densities were never more than 32 females/ha, but remained higher during winter. The spatial spread of the species varied between a few hundred meters to 2 km/year and was affected by the structure of the road network, topography and urban sprawl along the coast in the surrounding of the port of introduction. The study areas of Genoa, Venice and Rotterdam were found not suitable for establishment of this mosquito species, however climate change could create conditions for *Ae. aegypti* invasion in these regions in the next decades.

It is commonly accepted that targeted monitoring and early control actions are the most effective methods to hinder the establishment of invasive species in new areas. Our findings and model framework may support surveillance initiatives for those European coastal urban areas which have a known high propagule pressure and a high modelled probability of *Ae. aegypti* establishment.

**Highlights:**

- European coasts present favourable conditions for *Aedes aegypti* establishment
- We assess the species introduction and establishment using a process-based model
- We selected five ports: Algeciras, Barcelona, Venice, Genoa and Rotterdam
- Algeciras and Barcelona were the most suitable areas for the species establishment
- Climate change could make more suitable the northernmost study areas

## 1 Introduction

Mosquitoes in the genus *Aedes* (Culicidae: Diptera) are amongst the most feared invasive species due to their competence for transmitting debilitating pathogens. Among them, *Aedes aegypti* is a global health concern due to its capacity to thrive in urban areas and because it is an unrivalled vector for viruses (Leta et al., 2018). This species evolved in Sub-Saharan Africa and was progressively brought outside its original geographical range by human trades. First, the slave trade between the 15th and 19th centuries moved the mosquito from West Africa to Europe and the Americas. Afterwards, intensified trade with Asia during the 18th and 19th centuries and troops movements during World War II caused further expansion of its geographical distribution (Powell et al., 2013).

To date, mainland Europe is the only continent, except Antarctica, where *Ae. aegypti* has no established populations, despite having been present from Portugal to the Black Sea until the 20th century (Cardamatis, 1929; Schaffner and Mathis, 2014). Improvements both in hygiene and in the management of water systems as well as the widespread application of insecticides to decrease the malaria burden are thought to have led the species to extinction in this continent. However, in recent years *Ae. aegypti* has been detected multiple times in mainland Europe or in areas geographically or politically connected with Europe. Multiple introductions of *Ae. aegypti* were detected in the Netherlands in 2010 and again in 2016 (Brown et al., 2011; Ibañez-Justicia et al., 2017). Established populations of *Ae. aegypti* were found along the coast of the Black Sea in 2008 (in Turkey, Georgia and Russian Federation; ECDC et al. 2019), in the Portuguese Island of Madeira, located about 900 km South-West of mainland Portugal, in 2004 and, more recently, from nearby Egypt (Abozeid et al., 2018).

Mainland Europe has not yet witnessed the establishment and diffusion of *Ae. aegypti* populations. However, urban areas distributed along coastlines represent suitable habitats for the species (Kraemer et al., 2015). European coastlines are also characterized by extremely high exotic species propagule pressure (Lockwood et al., 2005; Dunn and Hatcher, 2015; Blackburn et al., 2020), due to the dense international maritime trade which nowadays accounts for 90% of the global exchange of goods in Europe (IMO, 2019). The Mediterranean basin plays a relevant role in global trades and it is expected to become even more important in the next decades since massive investments are planned in coastal and harbour infrastructures (e.g. see for instance the Chinese government’s Belt and Roads initiative; Ekman, 2018). Moreover, the Near and the Far East as well as North America, where *Ae. aegypti* populations are widespread, represent the major sources of goods shipped to Europe. Therefore, in the European context, coastal urban areas and the Mediterranean basin should be considered as focal points when evaluating the potential introduction of invasive mosquitoes. The precedent provided by *Ae. albopictus*, an invasive species related to *Ae. aegypti* could serve as a cautionary tale. This species was detected for the first time in the port of Durazzo, Albania, likely arrived from China (Adhami and Reiter, 1998), and nowadays is distributed along most of south and central Europe. Its introduction, establishment and dispersion in the European continent appear to have been supported by maritime transportation of tires and other goods (Eritja et al., 2005).

Outbreaks caused by *Aedes*-borne pathogens such as chikungunya and dengue have been recorded multiple times in Europe in the last two decades and, more recently, the first cases of Zika infection in humans have been reported in Southern France (Rezza et al., 2007; Venturi et al., 2017; Brady and Hay, 2019). Until now, these outbreaks have been limited in time and space and sustained by *Ae. albopictus*, which is an abundant but more opportunistic feeder and, thus, less competent vector than *Ae. aegypti* (Richards et al., 2006). The establishment of *Ae. aegypti* in Europe would dramatically change the epidemiological landscape by increasing the risk of pathogen transmission among the human population by this species, mimicking the public-health emergency which followed its recent introduction in the West Coast of the US (Metzger et al., 2017).

Surveillance enabling early warning and rapid control actions is the best tool we have to prevent species invasions, eradicate invading species or manage their population at low levels (Simberloff, 2014). Here, to support surveillance and early detection of *Ae. aegypti* in Europe, we applied a process-based population model which simulates both life cycle and dispersal of *Ae. aegypti* after its introduction, for example through ship traffic in European ports. To develop the model, we followed the theoretical model structure proposed by (Otero et al., 2008) and extended in Montecino et al. (2016). We chose to simulate introductions of *Ae. aegypti* in ports located along a latitudinal gradient to capture the variability of climatic condition along European coastlines. We selected five of the busiest European container ports (Eurostat, 2018) which were also located in areas predicted to be suitable for *Ae. aegypti* (Kraemer et al., 2015, 2019) *or where Ae. aegypti* was detected in the past, namely: Algeciras and Barcelona, Spain; Venice and Genoa, Italy and Rotterdam, the Netherlands.

The main questions we asked in this study were:

- Could *Ae. aegypti* eggs introduced via ship traffic via the major European harbours establish viable populations?
- If so, what is the likelihood that an introduction event brings about established invasive populations?
- At what latitude is an introduction more likely to happen?
- What is the minimum propagule pressure required for a successful introduction?
- What is the space-time spatial distribution trend which *Ae. aegypti* populations will follow once established?

## 2 Methods

### 2.1 Overview of model structure

We applied a process-based and time-discrete model to simulate population dynamics of *Ae. aegypti* after introduction in five study areas. *Ae aegypti* life cycle presents four main developmental stages: egg, larva, pupa and adult stage. We simplified the life cycle of *Ae. aegypti* by merging the larval and pupal stages in a unique compartment which we called “immature”. Thus, the model considers three main compartments: egg, immature and adult. Temperature and space can be considered the main environmental drivers of invasive mosquito ecology, thus events in the mosquito life cycle were treated as stochastic processes with probabilities derived from temperature-dependent (development) or distance-dependent (movement) functions (Fig. 1). Each event in the simulated life cycle occurred once per day always in the same order and each study area was divided in a grid composed of 250 m cells, which represented the fundamental spatial units into which*Ae. aegypti* life cycle took place and among which adult mosquitoes dispersed. This simplified space-time arena allowed us to reduce dramatically the structure of *Ae. aegypti* life cycle, and therefore the computing burden of the model, while retaining temporal, spatial and “biological” resolutions relevant for our questions (Pascoe et al., 2019). Model output consisted in a numerical matrix containing the daily number of individuals in each life stage for each 250 m cell of each study area, for each iteration.

**Figure 1:**
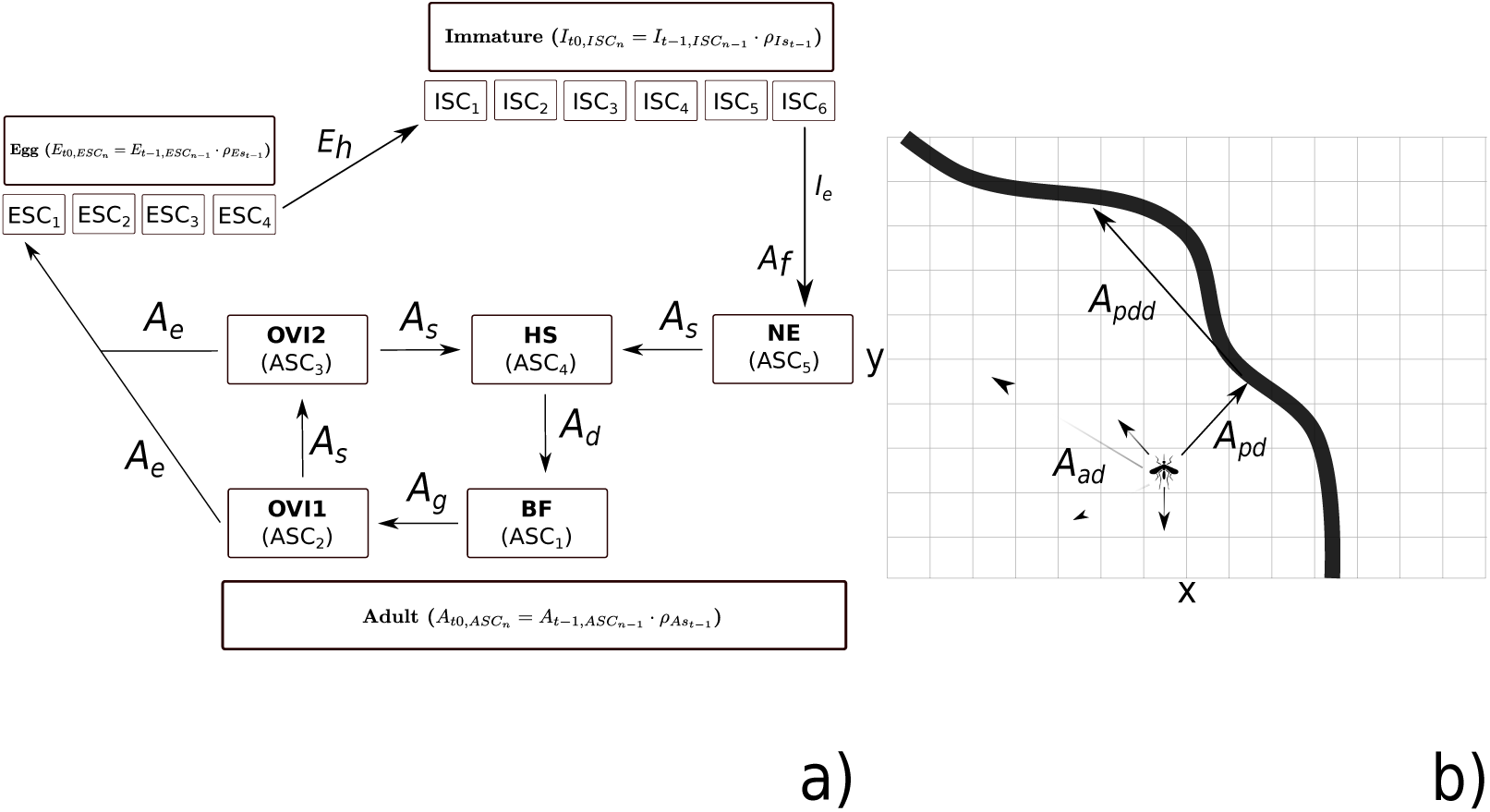
Graphical representation of model structure: in A) we showed the life cycle of *Ae. aegypti*, and in B) a representation of active and passive dispersal processes. A more detailed description of all model parameters reported in the figure is provided in Table 1.

### 2.2 Study areas and road network

We considered five 100×100 km study areas placed along a gradient of latitude (Fig. 2) and centered on the ports of Algeciras (lat=36.12, long=-5.43; Spain), Barcelona (41.34, 2.16; Spain), Genoa (44.40, 8.92; Italy), Venice (45.44, 12.25; Italy), and Rotterdam (51.90, 4.50; the Netherlands). These study areas have very different climatic conditions: Algeciras has a hot-summer Mediterranean climate with average annual temperature of 18.6°C and cumulative precipitation of 768 mm; in Algeciras summers are generally hot and dry, whereas winters are mild and wet. Barcelona and Genoa have a hot-summer Mediterranean climate but are surrounded by subtropical areas, reporting average temperatures of 21.2 and 16.7°C and cumulative precipitation of 640 and 1082 mm. Venice is characterized by a humid subtropical climate, with average annual temperature and cumulative precipitation of 14.5°C and 1101 mm, respectively. Winters in Venice can be cold, with temperatures commonly below 0°C during December and January. Finally, Rotterdam has a temperate oceanic climate, with average temperature and precipitation of 10.9°C and 867 mm, respectively. Rotterdam summers are generally cool and winter months rarely record temperature below 0°C (Fig. 3; Smith et al. (2011); Beck et al. (2018)). In addition to variability in the temperature regime, the five study areas show different environmental and topographic characteristics which represent natural barriers for species active dispersal. Such barriers can be overcome by invasive mosquitoes, nevertheless, through human-aided dispersal, with movements facilitated by human transportation (Service, 1997). The road network connecting European cities is amongst the most developed in the world, ensuring fast and easy movements of humans, goods and, unintentionally, introduced mosquito populations (Díaz-Nieto et al., 2016; Nelson et al., 2019). Here, in order to inform passive dispersal of *Ae. aegypti*, we considered only the network of primary roads in each 100 *km*^2^ study area (Center for International Earth Science Information Network - CIESIN - Columbia University and Information Technology Outreach Services - ITOS - University of Georgia, 2013). Moreover, to consider the eusynantropic ecology of *Ae. aegypti* (ECDC et al., 2019), we constrained both its life cycle and movements within urban areas, whose boundaries were defined using a spatial GIS layer derived from Schneider et al. 2009.

**Table 1:** A summary of model processes happening in day *t* and in cell *i*, with associated probability density distribution and parameters.

| Stage | Process description | Process ppd | Parameters | References |
| --- | --- | --- | --- | --- |
| Egg | **Survive:** Probability that an egg survives in day $t$, cell $i$. | $E_s \sim Uniform(p_{E_s})$ | $p_{E_s} = 0.99$ | Otero et al. (2008) |
| | **Hatch:** Probability that an embrionated egg hatches in day $t$, cell $i$. | $E_h \sim Binomial(p_{E_h})$ | $p_{E_h} = Beta(a, b);$<br>$a = 1.45, b = 6.51$ | Soares-Pinheiro et al. (2016) |
| Immature | **Survive:** Probability that an immature survives in day $t$, cell $i$. | $I_s \sim Binomial(p_{I_s})$ | $p_{I_s} = \mathcal{P}_6(temp)$ | Yang et al. (2009) |
| | **Emerge:** Probability that an immature $\geq 6$ days old emerges in day $t$, cell $i$. | $I_e \sim Binomial(p_{I_e})$ | $p_{I_e} = \mathcal{P}_7(temp)$ | Yang et al. (2009) |
| Adult | **Survive:** Probability that an adult survives in day $t$, cell $i$. | $A_s \sim Binomial(p_{A_s})$ | $p_{A_s} = \mathcal{P}_5(temp)$ | Yang et al. (2009) |
| | **Determine sex:** Probability that a newly emerged adult is female in day $t$, cell $i$. | $A_f \sim Binomial(p_{A_f})$ | $p_{A_f} = 0.5$ | - |
| | **Mature eggs:** Probability that a blood-fed female matures eggs in day $t$, cell $i$. | $A_{em} \sim Binomial(p_{A_{em}})$ | $p_{A_{em}} = \mathcal{P}_5(temp)$ | Yang et al. (2009) |
| | **Oviposit:** Number of eggs that an adult female, which underwent eggs maturation, lays in day $t$, cell $i$. | $A_{el} \sim Poisson(\lambda_{el})$ | $\lambda_{el} = \mathcal{P}_4(temp)$ | Yang et al. (2009); Christophers (1960b) |
| | **Disperse a-distance:** Probability that an host-seeking adult female disperses in day $t$ from cell $i$ to cell $i'$ at distance $d$. | $A_{ad} \sim Binomial(p_{A_{ad}})$ | $p_{A_{ad}} = LNormal(m, sd);$<br>$m = 4.95$<br>$sd = 0.66.$ | Marcantonio et al. (2019) |
| | **Disperse p:** Probability that an host-seeking adult female in day $t$ in cell $ii$ adjacent to a road intercepts a car. | $A_{pd} \sim Binomial(p_{A_{pd}})$ | $p_{A_{pd}} = 0.0051$ | Eritja et al. (2017) |
| | **Disperse p-distance:** Probability that a passively dispersing adult $A_{pd}$ disperses in day $t$ from cell $ii$ to cell $ii'$ at distance $d$. | $A_{pdd} \sim Binomial(p_{A_{pdd}})$ | $p_{A_{pdd}} = Gamma(\alpha, \beta)$<br>$\alpha = (c/10000)^2$<br>$alpha = c/10000^2$<br>$c = \{18.43, 23.14, 28.14\}$ | Alemanno et al. (2012) |

**Figure 2:**
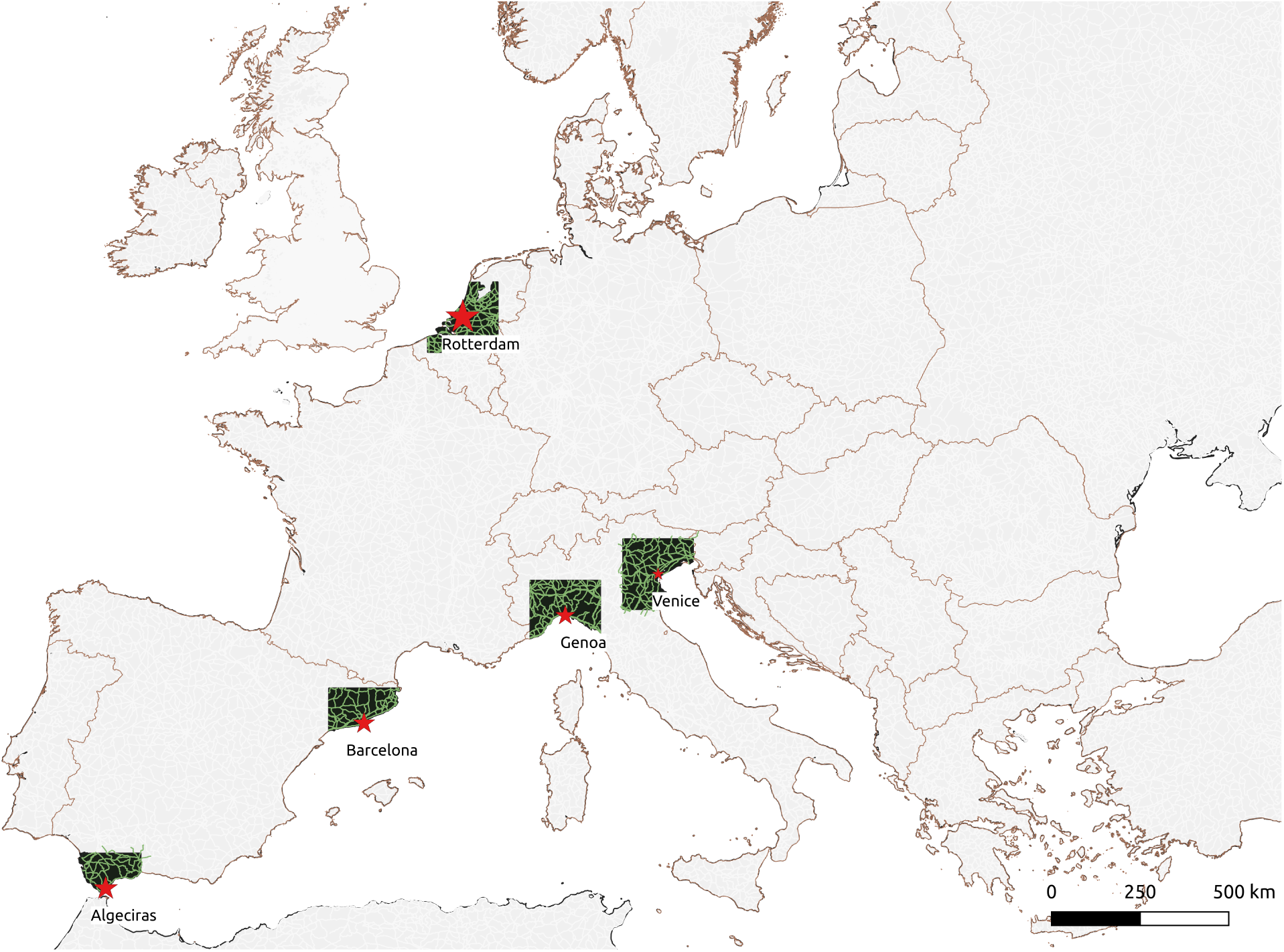
Locations of the five ports (red stars) investigated along a latitudinal gradient and corresponding study area (black). The network of primary roads is reported in green. The size of the “port” pin is proportional to the quantity of twenty-foot equivalent unit (TEU) received in the corresponding port for the year 2018 (Amsterdam=13.6, Algeciras=4.8, Barcelona=3.4, Genoa=2.5, Venice=0.6 TEU).

**Figure 3:**
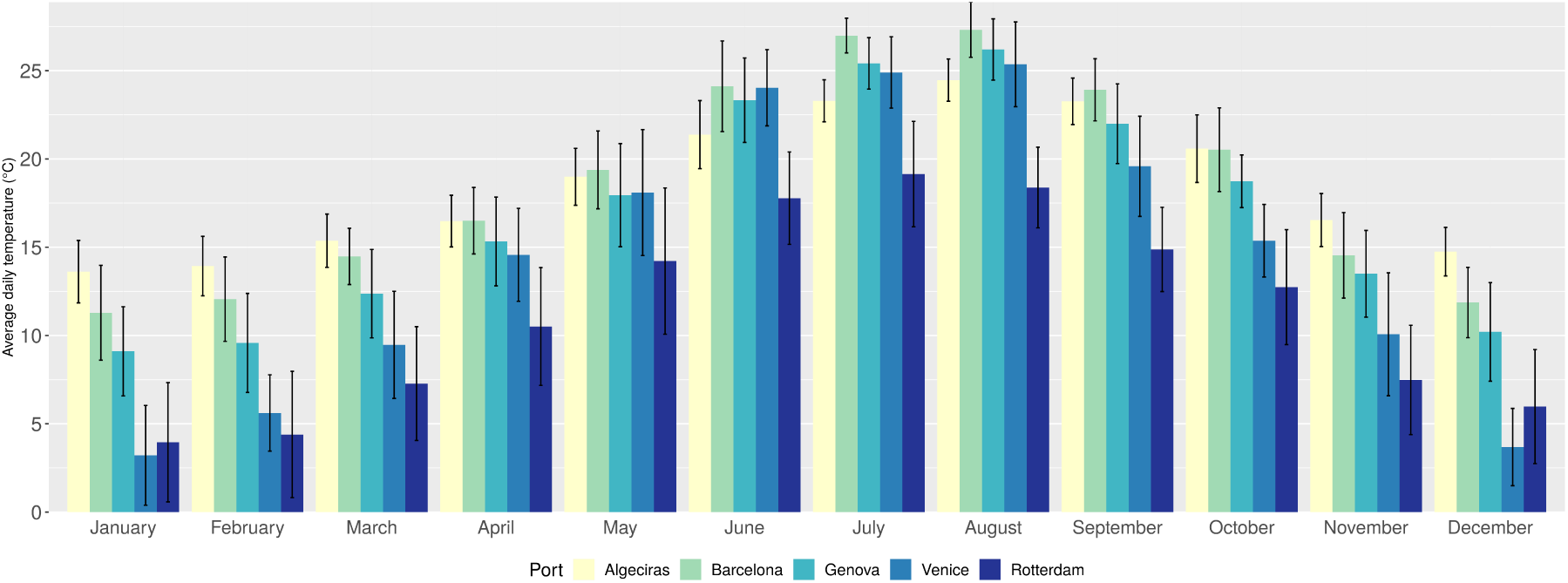
Monthly average temperature and standard deviation (error bars) for the years 2017-2019 in the five study areas.

### 2.3 Temperature data

Daily temperature estimates at 2 m above-ground for the period 2016-2019 were obtained using the R package *microclima* (Kearney et al., 2020) for each study area, considering a region spanning 100 *km*^2^ from the center of each port. Microclima makes use of 6-hour climate and radiation data downloaded from the National Center for Environmental Prediction Reanalysis 2 database at a resolution of 1.8-2.5° (Kanamitsu et al., 2002). The four daily observations are then interpolated hourly with spline interpolation and corrected considering topographic effect using a digital terrain model at 250 m resolution downloaded from EUDEM (Copernicus project, 2019). The microclima package allows to define an habitat category as defined in MODIS Land use datasets (MODIS product MCD12Q1v006) (M. Friedl, 2015), which is used to calibrate a Gaussian curve to estimate hourly vegetation features using a-priori established relationship between habitat type and observed vegetation characteristics. We decided to consider the vegetation cover across our study areas according to the MODIS habitat class closer to the ecology of *Ae. aegypti*, which is “Urban and built-up” (MODIS category 14). Coastal effects on temperatures were also considered as described in (Maclean et al., 2019). Finally, the temperature times series for each study area derived with Microclima was further fine-tuned to local weather conditions using data from the weather station from the NOAA network (Smith et al., 2011) closest to the relevant port.

### 2.4 Description of the population dynamical model

In the proposed model arena, which we described by space *i* and time *t*, the number of individuals in each developmental stage dying or surviving day *t* in cell *i* is an outcome of binomial draws with probabilities derived from temperature-dependent functions parameterized with data from the appropriate scientific literature. All processes with their associated probability distributions, parameters and references are described in Table 1.

The egg stage (E) is composed by four sub-compartments (ESC) which contain eggs of different ages. Eggs enter ESC-1 when laid by ovipositing adult females; the first three sub-compartments contain eggs one, two and three days old that are undergoing embrionation (i.e., incubation time; Christophers (1960a)). The number of eggs (*E*) in *sc*-1 during day *t-1* that die or move to *sc* in day *t* is defined by a binomial random draw based on mortality and survival probabilities 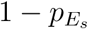 and 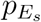, respectively:

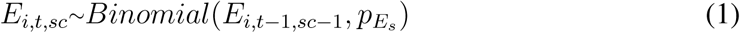

Four days-old eggs can hatch as immature to move to the “immature” (*I*) compartment. The hatching event occurs with probability *p*_*Eh*_, which is established by a random draw from a Beta distribution. The two shape parameters of this Beta distribution were derived using the average (0.076) and low 95% CI (0.023) proportion of *Ae. aegypti* eggs hatched after being submerged in plastic containers filled with water in field conditions (SoaresPinheiro et al., 2016).

The immature compartment is composed by six sub-compartments (ISC) that contain immatures of different age or stage. Every day immatures in ISC-1 to ISC-5 can die or move to the next ISC based on temperature-dependent probability 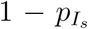 and 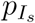, respectively.

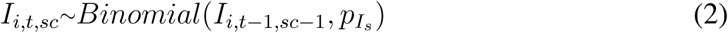

When immatures reach ISC-6, they become ready to emerge as adults and therefore can die, survive as other ISC-6, or emerge to move to the “adult” compartment with probability 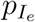, which is the outcome of a temperature-dependent polynomial function as defined in (Yang et al., 2009). The pool of immatures moving every day to the adult compartment is subjected to an additional Binomial random draw in order to limit the adult compartment to only female mosquitoes (assuming a 1:1 male/female ratio).

The adult compartment is composed by five sub-compartments that contain adult females in different ages or physiological status. ASC-1 contains 1-day old adults which are developing gonads and therefore are not yet sexually mature (Christophers, 1960a). Adults in ASC-1 can die or move to ASC-2, which contains host-seeking adults, following a random draw with temperature-dependent probability 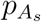 (Yang et al., 2009), which also drives the fate of ASC-2 to 5. ASC-2 can die or move to ASC-3 where they are assumed to have taken a blood meal and entered the gonotrophic cycle. ASC-3 is therefore composed of blood-fed adult females maturing eggs (i.e., completing the gonotrophic cycle). If adult females in ASC-3 survive day *t-1* they can complete the gonotrophic cycle and move in day *t* to ASC-4 where they become ovipositing females. The completion of the gonotrophic cycle by ASC-3 is driven by a binomial random draw based on temperature-dependent probability 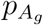 (Yang et al., 2009). Females in ASC-4 in day *t*, first oviposit, then die or move to ASC-5 which contains females in the second day of oviposition. The number of eggs laid by a female in ASC-4 or ASC-5 is the outcome of a Poisson random draw with temperature-dependent probability 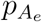 (Yang et al., 2009; Christophers, 1960a). Finally, females in ASC-5 die or move to ASC-2, where they will seek again for a blood-meal and, if surviving, re-enter the gonotrophic cycle as described above.

In addition to the development and survival cycle of events, host-seeking adult females (ASC-4) can also disperse actively (short-range dispersal) or passively (medium-range dispersal). In our model, active dispersal is defined as the probability that an adult mosquito flies at distance *d*_*ad*_ from the cell of origin and follows a log-normal dispersal kernel (Marcantonio et al., 2019). Thus, in each day *t*, the number of dispersing ASC-4 that move to a distance *d* from the origin cell *i* is an outcome of a binomial random draw with probability 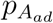:

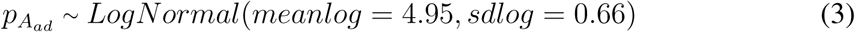

We assumed that females start dispersing from the center of the cell of origin *i*, therefore, given a cell size of 250 m, all mosquitoes dispersing less than 250 m do not leave *i*. All other dispersing mosquitoes move to a cell *i*′ (landing cell) which is chosen randomly from the set of cells at distance *d* (i.e. non directional active dispersal).

In addition to active dispersal, the proposed model also considers dispersal aided by car traffic along the main road network. This type of dispersal is thought to be amongst the main drivers of medium-range geographical expansion for *Aedes* mosquitoes (Marcantonio et al., 2016; Eritja et al., 2017). In the model, adults of all ages, except 1-day old, that reside in cells intersecting roads may undergo this type of dispersal. We parameterized passive dispersal using the probability of a car transporting *Aedes* mosquitoes 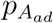 of 0.0051, as reported in Eritja et al. (2017), implying that, on average, only about 5 over 1,000 mosquitoes disperse passively every day from each cell of origin. Furthermore, to define the probability of dispersing passively at different distances *d*, we used data on driving patterns in Europe (Alemanno et al., 2012). We defined 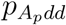 as the outcome of a Gamma distribution with mean, *md*, equal to the average driving distance per trip in each of the study areas:

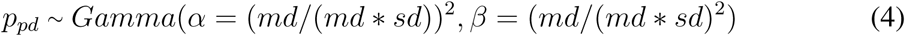

where *md* was set to 18.43 km for Italy, 23.14 km for Spain and 28.14 km for the Netherlands (Alemanno et al., 2012). The standard deviation used to parameterize the Gamma distribution was chosen to be very large (100 times the mean) in order to assign a small probability density also to large dispersal distances. The matrix of distances between cells along roads was calculated using the set of *v.net* modules in GRASS GIS 7.6 (Neteler et al., 2012).

The model coded in the R statistical language (R Core Team, 2019) and adapted for parallel computation is available in the Github repository: https://github.com/mattmar/euaeae.

### 2.5 Experimental design and model validation

We simulated the introduction of 10, 50, 100, 250, 500 or 1000 eggs in a randomly chosen cell randomly within the boundary of the corresponding port in each study area. Eggs were chosen as the introduction propagule as they are the resistant life stage through which *Aedes* mosquitoes overcome periods of adverse climatic conditions, such as cold winters or dry summers. Each introduction was iterated 500 times to account for stochastic events in both life cycle and dispersal, as the coefficient of variation of all outcomes of interest stabilised around this number of iterations. The length of each simulated introduction was set to three years in order to allow stabilisation of the simulated populations and integrate inter-annual climatic variability. The introduction year was 2017. In addition, we decided to run preliminary simulations to define the most favourable period of the year for introducing *Ae. aegypti* in order to reduce the chance of false negative results. This set of simulations began with 100 adults introduced on the 15th of each month of the year 2017 and lasted until the end of the same year. The introduction date in 2017 was thus chosen for each study area as the starting date of the scenario that recorded the highest number of adult mosquitoes in any of the simulated days. In the subsequent full model simulations, successful introductions were defined as those that yielded at least one individual in any life stage on the 20th of June 2018, which represents the end of the spring (and therefore the end of adverse climatic conditions for *Aedes aegypti*) the year after date of introduction (2017). Population demographic and dispersal trends for each introduction scenario were summarised using the inter-quartile ranges of the distribution of each outcome of interest across iterations for each simulated day.

To validate model outputs, we derived the proportion of each study area which experienced days below the 10°C isotherm for the whole 2017-2018 period, using ERA5 re-analysis (1979–2018) (Copernicus Climate Change Service, 2017). Temperatures around 10°C represent a theoretical thermal limit for *Ae. aegypti*, below which adult mosquitoes become torpid and unable to move (Christophers, 1960a; Reinhold et al., 2018). The invasion success ratio of simulated introductions was compared against the proportion of the study area below the 10°C isotherm to assess the credibility of model outputs.

Local sensitivity analysis was performed on model results by varying the values of four model parameters which were considered uncertain, namely: the probability of egg survival 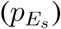, the probability of egg hatching 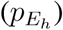, the number of introduced eggs (*E*), the probability of adult dispersing passively 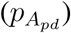. We performed the sensitivity analysis by running 200 model iterations over an extensive range of values of each selected model parameter for the study area of Algeciras. The proportion of successful iterations, maximum population dispersal distance, female abundance and invaded area in hectares were plotted against each parameter range in order to assess the sensitivity of model outputs. Results of the local sensitivity analysis are reported in Appendix 2.

## 3 Results

### 3.1 Likelihood of a successful introduction relative to time of the year

Results from the set of preliminary model simulations showed that the month of introduction which resulted in the highest median abundance of adult mosquitoes (relatively to the first year of introduction) was March for the port of Rotterdam, April for Barcelona, and May for the ports of Genoa, Venice and Algeciras. Introductions in any month of the year 2017 showed viable populations on the last day of the year (i.e., successful introductions) only in Barcelona, while Algeciras (10 months), Genoa (9), Venice (3) and Rotterdam (1) showed a decreasingly lower number of months suitable for successful introductions (Fig. 4).

**Figure 4:**
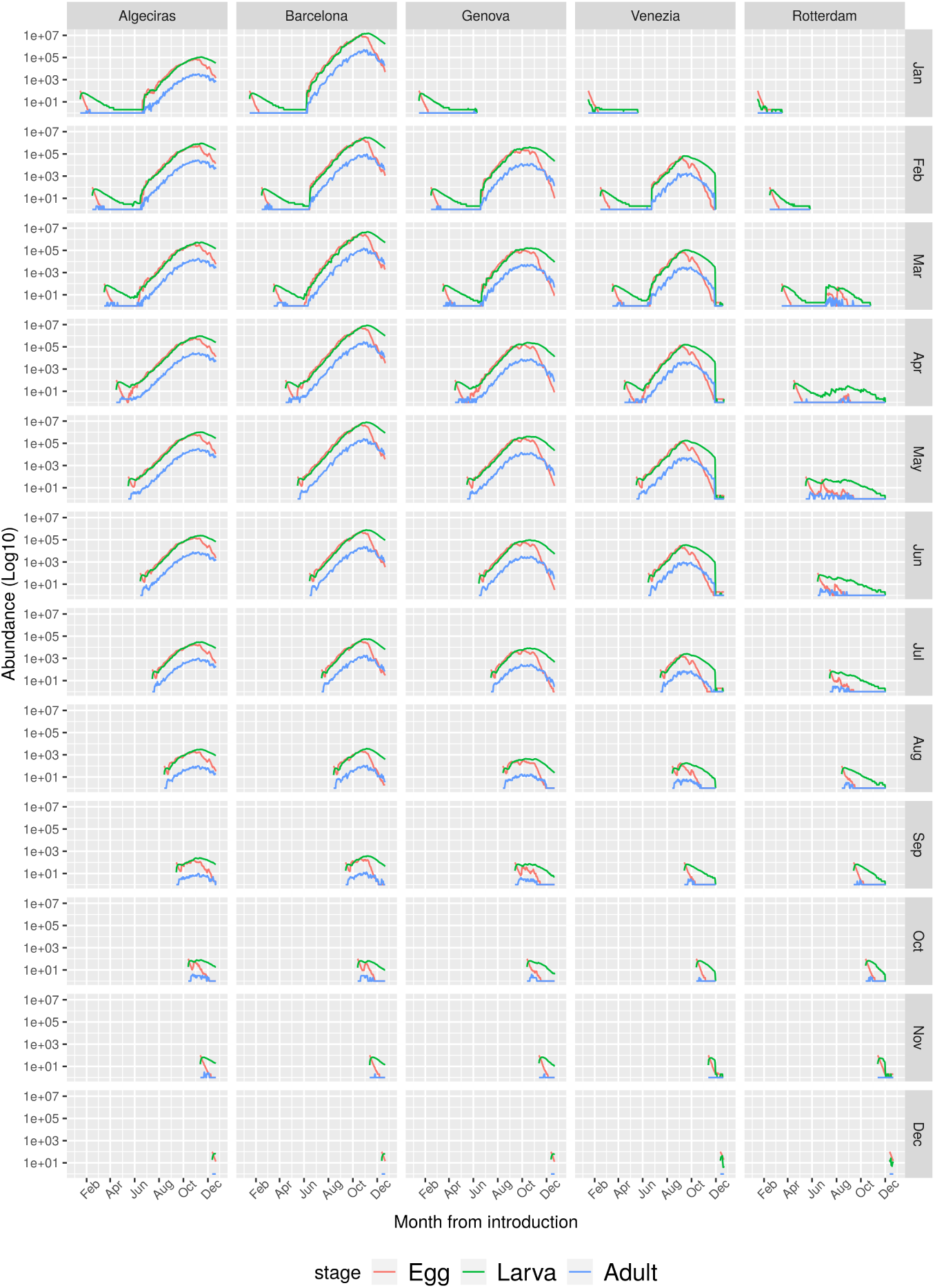
Time trend of *Ae. aegypti* population abundances from preliminary model iterations simulating the introduction of 250 eggs starting on the 15th of each month of 2017 and ending on the 20th of December 2017 for the five study areas. Blue lines represent adult females, green lines larvae and red lines eggs. Values on the y-axis are transformed using the common logarithm.

### 3.2 Likelihood of a successful introduction relative to latitude

The percentage of full model simulations which resulted in established *Ae. aegypti* population was 100% for Barcelona, for a propagule pressure of at least 250 eggs. Algeciras showed somewhat similar results, however, 100% successful introductions was achieved only with a higher minimum propagule pressure of 1000 eggs (Table 2). On the contrary, we did not observe successful introductions for any propagule pressure either in Genoa, Venice or Rotterdam. In Rotterdam, mean daily temperatures during summer were never high enough (>20°C) to allow the accumulation of an adequate egg bank which would have permitted the overwintering of the population (see Fig. 3). Conversely, in Genoa and Venice, although introductions sometimes caused moderate population abundances (>50 adult mosquitoes per ha) during the most favourable months, very cold winter temperatures, often under 0°C, caused extinction for all the introduced populations. The percent success of invasion was well correlated with the percentage of area under the 10°C for the period 2017-2018. Interestingly, a relatively small decrease of about 6% of area-days under the 10°C isotherm between the ports of Barcelona and Genoa represented a climatic divide for *Ae. aegypti* invasion success.

**Table 2:** Number of introduced eggs and corresponding percentage of successful introductions for each of the considered study areas. The last column represent the proportion of each study area that fell below the isotherm of 10°C for the years 2017-2018 (ERA5 reanalysis (1979–2018)).

| Site | Introduced eggs | Introduction success % | Proportion of the study area below 10°C % |
| --- | --- | --- | --- |
| *Algeciras* | 10,50,100,250,500,1000 | 6,11,16,48,74,100 | 4.8 |
| *Barcelona* | 10,50,100,250,500,1000 | 4,26,72,100,100,100 | 15.3 |
| *Genoa* | 10,50,100,250,500,1000 | 0 | 21.7 |
| *Venice* | 10,50,100,250,500,1000 | 0 | 24.6 |
| *Rotterdam* | 10,50,100,250,500,1000 | 0 | 29.4 |

### 3.3 Space-time trend of introduced populations

With the exception of Rotterdam, all simulated populations were able to reproduce and spread at local scale at least during the first summer after introduction. The peak of adult mosquito abundance was reached in autumn in Algeciras, Barcelona and Genoa or in late summer in Venice. Adult female abundances reached maximum overall values of 584 *· ha*^−1^ in Barcelona, but were considerably lower in the other study areas: Algeciras yielded a maximum of 32 *· ha*^−1^, Genoa 28 *· ha*^−1^, Venice 20 *· ha*^−1^ and Rotterdam 2 *· ha*^−1^ (Fig. 5). The pools of eggs and immatures followed trends similar to adult mosquitoes but lagging in time proportionally to differential developmental times, and reaching higher abundances due to the inter-stage survival bottlenecks (Appendix Fig. 7). However, with the onset of colder weather conditions at the end of summer, mosquito populations declined rapidly in all the study areas. The decline was more evident as soon as daily average temperature was constantly under 10°C, and even in Algeciras and Barcelona, winter densities dropped to a minimum to 1 female/ha during winter months. In these months, the consistent egg bank accumulated during the warmer months ensured population overwintering. On the contrary, after the first summer, simulated populations from Genoa, Venice and Rotterdam were not able to overwinter (Fig. 5). The 10°C isotherm computed for the whole Europe from the first of September 2017 to the 31th of May 2018 showed that these three study areas, but not Algeciras and Barcelona, presented mean temperatures values below 10°C for at least 1 full winter month.

**Figure 5:**
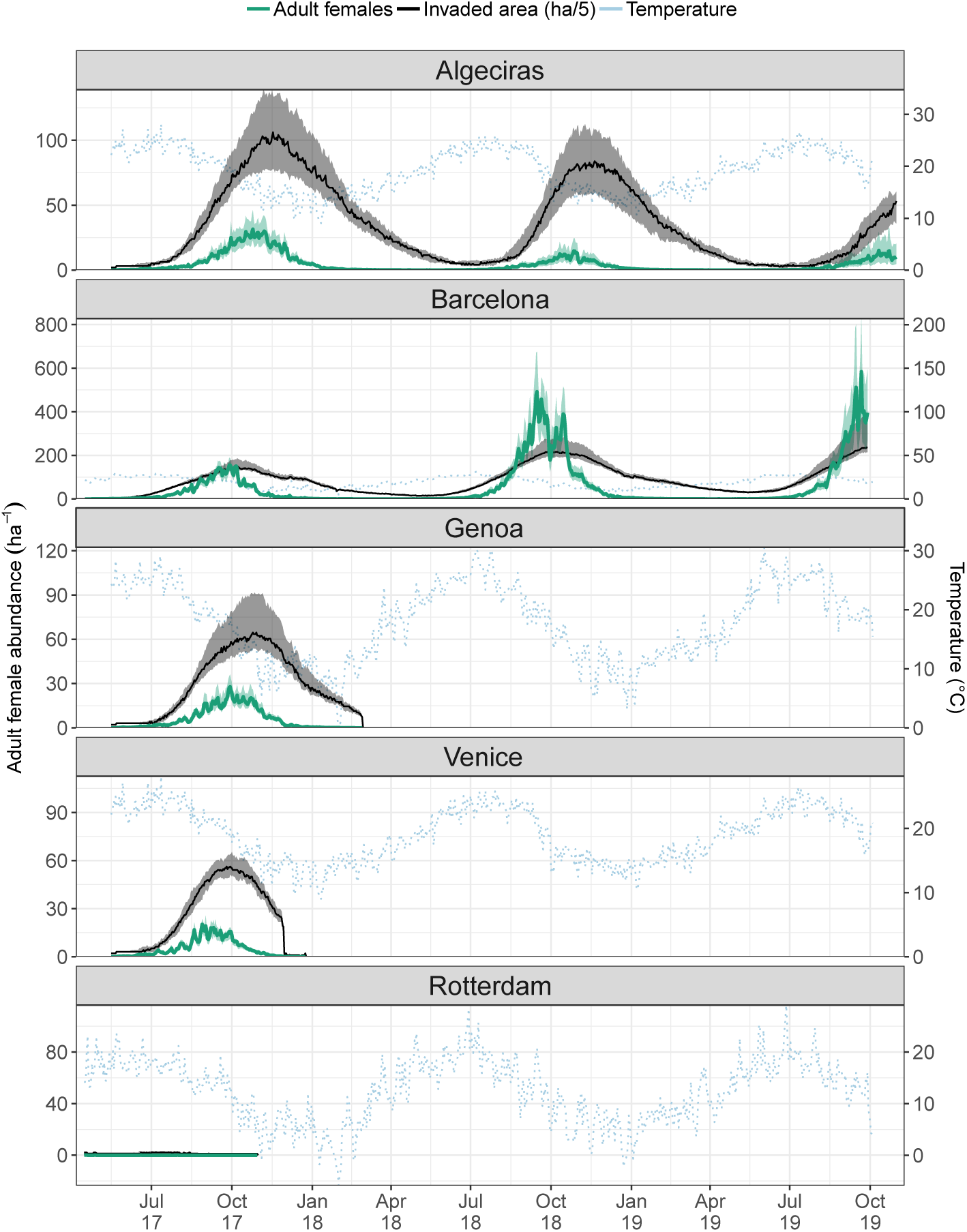
Temporal trend describing the outputs of model simulations for the five study areas. We reported both median (green lines) and inter-quartile (green ribbons) population densities of adult females per hectare and median (black line) and inter-quartile range (grey ribbons) of the invaded area in hectares (divided by a factor of 5 for scale reasons), derived from 500 iterated introductions of 250 *Ae. aegypti* eggs in each of the 5 study areas. The light-blue dotted line reports the trend in average daily temperatures and refers to the values in the right y-axis.

**Figure 6:**
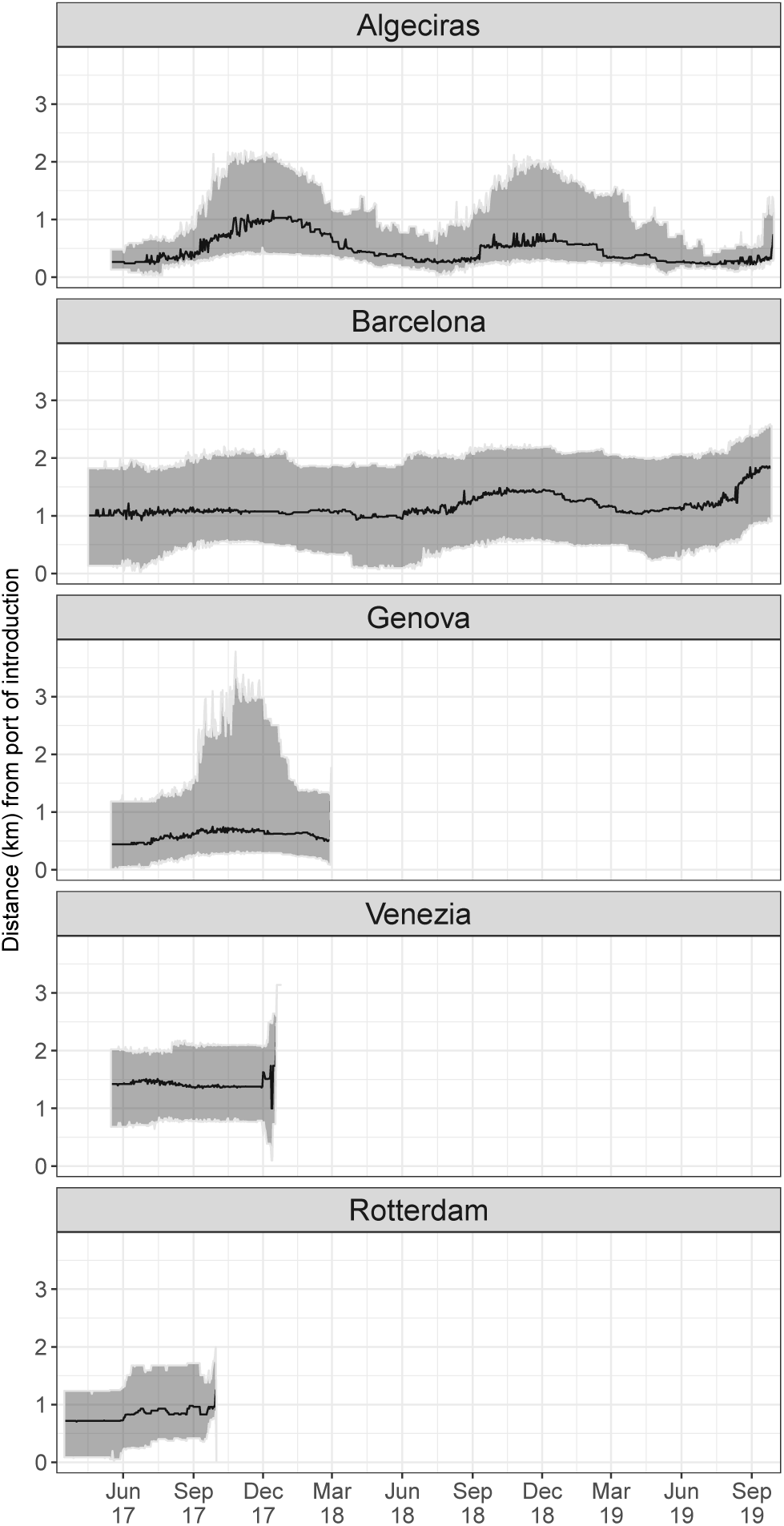
Median (black line) and inter-quartile (grey ribbons) dispersal distance (km) of simulated mosquito populations derived from 500 iterated introductions of 250 *Ae. aegypti* eggs in each of the 5 study areas. Dispersal distance was calculated as the euclidean distance between the centroid of each port of introduction and centroids of cells with at least one *Ae. aegypti* individual in any developmental stage.

The spatial spread of introduced populations was limited during the simulated period, as the median dispersal distance was never higher than 4 km from the port of introduction (Fig. 5). The two study areas which showed viable populations after the first simulated year reported different dispersal trends. *Ae. aegypti* populations in Algeciras showed a more seasonal spatial spread, dispersing more during summer and contracting during winter. The simulated populations in this study area moved farther just after introduction, dispersing more than 2 km and invading a maximum of 638 hectares, but in the following years the invaded area contracted. On the contrary, Barcelona’s populations remained in the surrounding of the introduction location for more than one year before showing a more marked spread. During 2019, this population was able to spread to a maximum of 2434 ha, doubling the area invaded in 2018 (Fig. 5).

## 4 Discussion

The overarching aim of this study was to examine both the likelihood and the dynamics of a putative re-introduction of *Ae. aegypti* in European ports under current climatic conditions. To do so, we developed a spatially explicit process-based model tailored to the physiological requirements of *Ae. aegypti*. We applied the model to simulate and study the introduction of *Ae. aegypty* in five major European ports with high potential propagule pressure and along a gradient of latitude. Our modelling framework accounted for: i) the effect of spatial and temporal temperature variation on population dynamics, ii) the dispersal at local and regional scales as well as iii) stochastic events following introduction. The approach we adopted has the advantage to be mechanistically linked to *Ae. aegypti* biology which stems in more ecologically-realistic estimates of the likelihood of invasion in already sensitive areas.

According to model results and given current climatic conditions, a relatively low *Ae. aegypti* propagule pressure has the potential to cause species establishment, high local densities and slow initial dispersal in the two southernmost study areas: Algeciras and Barcelona. Overall, mosquito densities reported in model outputs, with peaks of 584 females per hectare in Barcelona study area, are in line with previous observations in areas where *Ae. aegypti* is well established (Focks et al., 1981; Ritchie et al., 2013; Garcia et al., 2016).

Barcelona was the most suitable to invasive *Ae. aegypti* than Algeciras, allowing also for higher abundance and more rapid spatial spread of simulated invasive populations. These results are associated with the different layout of urban areas around the two ports as well as local climatic conditions. Algeciras is relatively isolated from other urban areas, therefore dispersing mosquitoes need to disperse relatively far to find suitable urban environments, which may have favoured a more seasonal dispersal pattern. This pattern may have only rarely allowed for enough propagules to “jump” to and colonise new distant areas, resulting in local extinction and invaded area contraction during winter months (Roche et al., 2015). On the contrary, the massive urban sprawl of Barcelona seamlessly connects the port area with neighbouring suitable urban areas along the coast either North, West or South (Fig. 5). As a result, invasive propagules could gradually form new invasion fronts, favouring a longer and more gradual dispersal. Both Algeciras and Barcelona fall in the hot-mediterranean climatic zone, which is characterised by warm summer and mild winters, meaning optimal conditions for *Ae. aegypti* life cycle. The relatively higher invasion success and population densities simulated for Barcelona can be explained considering this area warmer temperatures between May and September, which also correspond to the core of the mosquito growing season. Although we did not consider water availability for breeding sites in our model, it is worth highlighting that this would not represent a limiting condition in both the suitable study areas, as yearly precipitations are well above values of 400-500 mm/yr known to hinder *Aedes* population persistence (Caminade et al., 2012). Mosquito populations introduced either in the ports of Genoa, Venice and Rotterdam, despite persisting for short-to-longer periods and dispersing in the surrounding areas of each ports, were never able to establish overwintering populations. These three areas have longer and harsher cold temperature during winter months which are at least for one consecutive month below the isotherm of 10°C, making it impossible for introduced *Ae. aegypti* to overcome the adverse period of the year.

Taken together these results indicate that, regardless of the propagule pressure, the likelihood of *Ae. aegypti* invasion along European coasts is high in the southern Mediterranean basin, is less likely at latitudes between the ports of Barcelona and Venice, and becomes extremely unlikely further up North. However, where the current temperature regime is too cold to allow species survival over winter, either microclimatic refugia under current climatic conditions or climate change, in the near future, could still allow *Ae. aegypti* establishment. Microclimatic refugia are widespread in urban areas and can be caused by urban heat-island effects or by urban infrastructures and have been shown to be exploited by *Ae. aegypti* (Tsunoda et al., 2014). *An example of local conditions detached from the regional climatic regime which has allowed Ae. aegypti* persistence is the city of Washington, DC, USA, where *Ae. aegypti* was able to establish viable populations by overcoming the otherwise unsuitable climatic conditions sheltering in the myriad of underground refugia of the metropolitan area (Gloria-Soria et al., 2018).

Microclimatic urban refugia are by definition limited to small extensions, whereas global change has the potential to expand dramatically the geographical extent suitable for *Ae. aegypti* as well as *Aedes*-borne virus transmission in Europe. Mosquito physiological rates are shortened by increasing environmental temperatures in the range of 10-35°C (Mordecai et al., 2019), and the 10°C isotherm of the coldest month is often used as a proxy to assess *Ae. aegypti* geographic suitability. Europe is currently characterized by a 10°C isotherm of the coldest month extending to most of the south inland areas of Spain, Italy and Greece, which underpins how three out of five coastal areas that we tested were not suitable for *Ae. aegypti* establishment. However, the current direction of global change is predicted to bring warmer temperatures in most of Europe that, coupled with unchanged or increased precipitation along coastal areas, will impact the extent of regions where *Ae. aegypti* introduction could cause invasions (Kraemer et al., 2019; Liu et al., 2019; Ryan et al., 2019). That said, our findings suggest that simulated populations introduced in the region of Genoa persisted for most of the winter to became extinct only at the beginning of spring. Some model iterations showed individuals persisting till the month of April 2018. This result suggests that, according to our model, Genoa is close to suitability for *Ae. aegypti*. Thus, areas such as the region of Venice and Genoa, which have already acted as ports of introduction for *Ae. albopictus* (Sabatini et al., 1990; Knudsen et al., 1996), another invasive mosquito, should already be targeted for limiting the likelihood of *Ae. aegypti* establishment in the near future.

The potential establishment of populations of *Ae. aegypti* in the southernmost study areas considered in this study does not consequently imply an arbovirus outbreak. Howrever, assuming a putative introduction of *Aedes*-borne pathogens and the lack of adequate surveillance, the socio-environmental conditions of Algeciras and Barcelona may be suitable to sustain transmission of such introduced pathogens in the human population. In the past twenty years, the chikungunya outbreaks in southern european countries such as Italy, Spain and Croatia results from the establishment of *Ae. albopictus* in highly dense populated areas coupled with the introduction of pathogens via infected travellers returning from areas with high incidence rates (Liumbruno et al., 2008; Lindh et al., 2019).

Our modelling exercise is not exempt from caveats. We did not consider evolutionary processes which may affect mosquito physiological rates allowing *Ae. aegypti* to be locally adapted to different set of environmental conditions (Reinhold et al., 2018). For example, variability in the quiescence strategy to prolong embryonic viability has been observed in different *Ae. aegypti* populations (Oliva et al., 2018). Moreover, potential interspecific competition between individuals of *Ae. albopictus* and *Ae. aegypti* is another factor not considered in this study which may limit the invasion success of *Ae. aegypti* (Carrasquilla and Lounibos, 2015). Finally, we assumed that both human host density and the number of breeding sites are not limiting factors. We believe these assumption realistic, because: 1) we selected main European ports surrounded by urbanized areas as introducing sites, which are expected to have a high density of available hosts; 2) due to the domestic behaviour *of Ae. aegypty*; it is well know that *Ae. aegypti* has preference for man-made containers as breeding sites, and discarded objects such as cans, jars, pots are expected to be overabundant in urban areas. We also acknowledge that we have considered urban areas as homogeneous land use features, which is far from reality. Urban areas are dynamical landscape features, whose heterogeneity has been recognized as an emerging characteristic in pathogen circulation dynamics. The importance of urban areas for vector species lays in the very diverse microclimatic and habitat conditions they can provide, which creates emerging interfaces for interaction between humans and other species (Vanwambeke et al., 2019).

## 5 Conclusions

Environmental conditions in Southern European urban areas such as Algeciras and Barcelona suggest potential for colonization of these areas by *Ae. aegypti*. We found that the spread pattern in time and space following establishment could be a continuous gradual spread or a more seasonal stepping-stone pattern, dictated by the specific characteristics and connectivity of each urban areas. Moreover, in case of establishment, *Ae. aegypti* populations in these areas were found to reach abundances similar to regions of the world with active transmission of *Aedes*-borne viruses. European areas located at higher latitudes, such as the port of Genoa, were found not suitable for colonization by these mosquitoes, but Global Change could modify conditions for *Ae. aegypti* invasion also in these regions. Finally, colonization of Northern European urban areas, aside short-lived and limited encroachment due to urban refugia, was found to be very unlikely.

It is commonly accepted that targeted monitoring of sensitive areas and early control actions are the most effective methods to prevent the establishment of invasive species in new areas. Maintaining *Ae. aegypti* populations at low densities and possibly limiting their spatial spread is crucial to avoid the risk of pathogen transmission. Our findings and model framework may support surveillance initiatives for those European coastal urban areas which have a known high propagule pressure and a high probability of *Ae. aegypti* establishment.

The model we applied for our simulations is made available as an R function, in the hope to foster reproducibility, further development and application onto different scenarios, geographical areas or spatial scales.

## 6 Acknowledgements

DDR is supported by the FRS-FNRS.

## Appendix 1: Temporal and spatial trends of the simulated mosquito populations

### Temporal trend for all life stages

**Figure 7:**
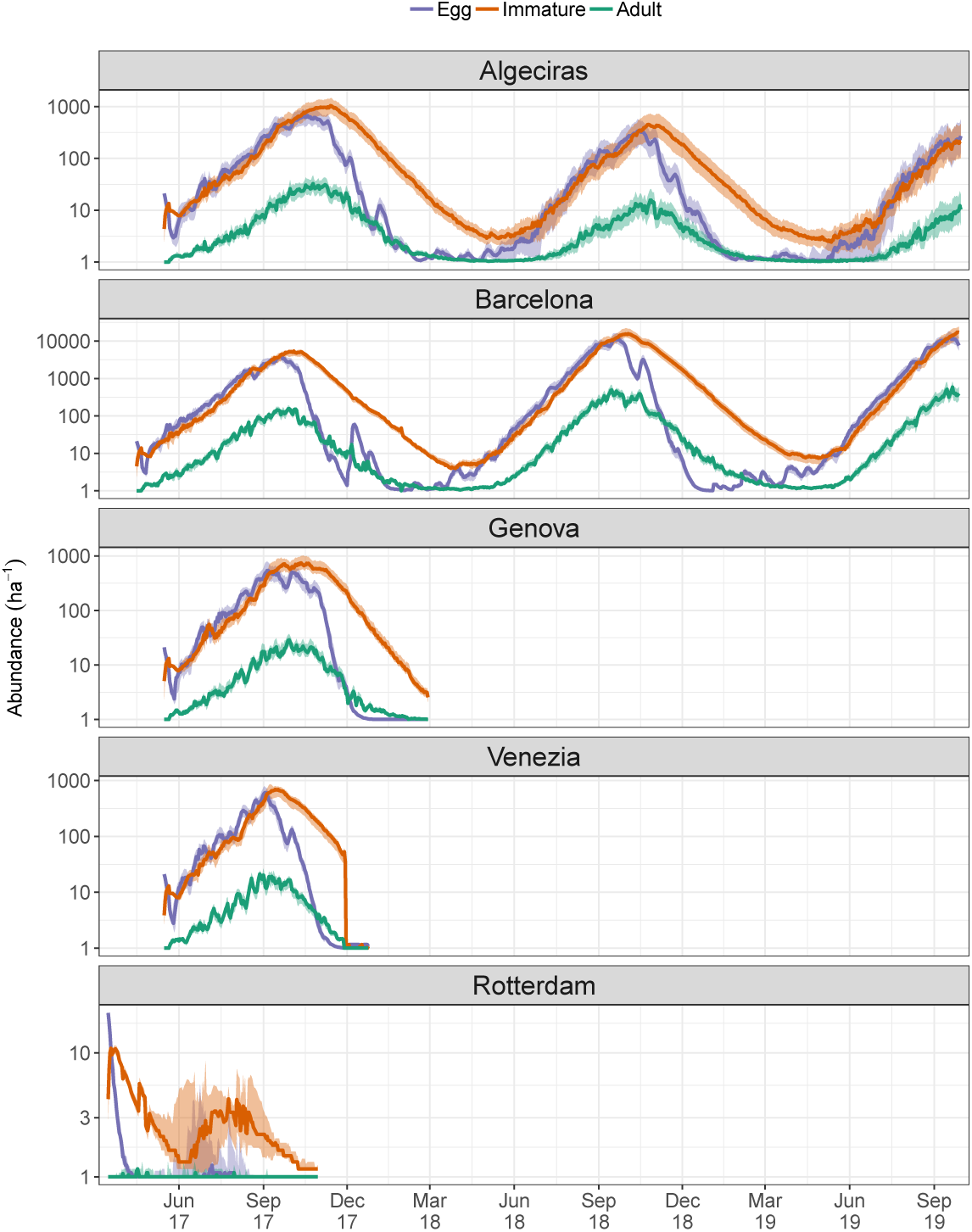
Temporal trend reporting: i) median (lines) and inter-quartile (ribbons) population abundances for each stage (adults in green, immatures in orange and eggs in purple) for introduced *Ae. aegypti* populations. Values on the y-axis are transformed using the common logarithm.

## Appendix 2: Sensitivity analysis on uncertain model parameters

In an attempt to assess and report the effect on model result of those model parameters which were judged more uncertain (i.e., lacking experimental or field estimates) we run a global sensitivity analysis by exploring the parameter space of: the probability of egg survival 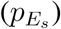, the probability of egg hatching 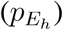, the number of introduced eggs (*E*), the probability of adult dispersing passively (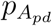; Table 3). We performed the sensitivity analysis iterating 200 times the introduction of 250 eggs (except for *E*) on the 15th of May on the study area of Algeciras.

**Table 3.**

| Model parameter | Parameter space | Value used in the model |
| --- | --- | --- |
| Daily probability of egg survival | (0.90,1.00) | 0.99 |
| Daily probability of egg hatching | (0.01, 1.00) | 0.076 |
| Daily probability of passive dispersal | (0.0018, 0.0180) | 0.0051 |
| Number of introduced eggs | (10,1000) | 250 |

Summary statistics of model results were plotted against the range of parameter values to provide a simple and straightforward assessment of model sensitivity (Fig. 8).

**Figure 8:**
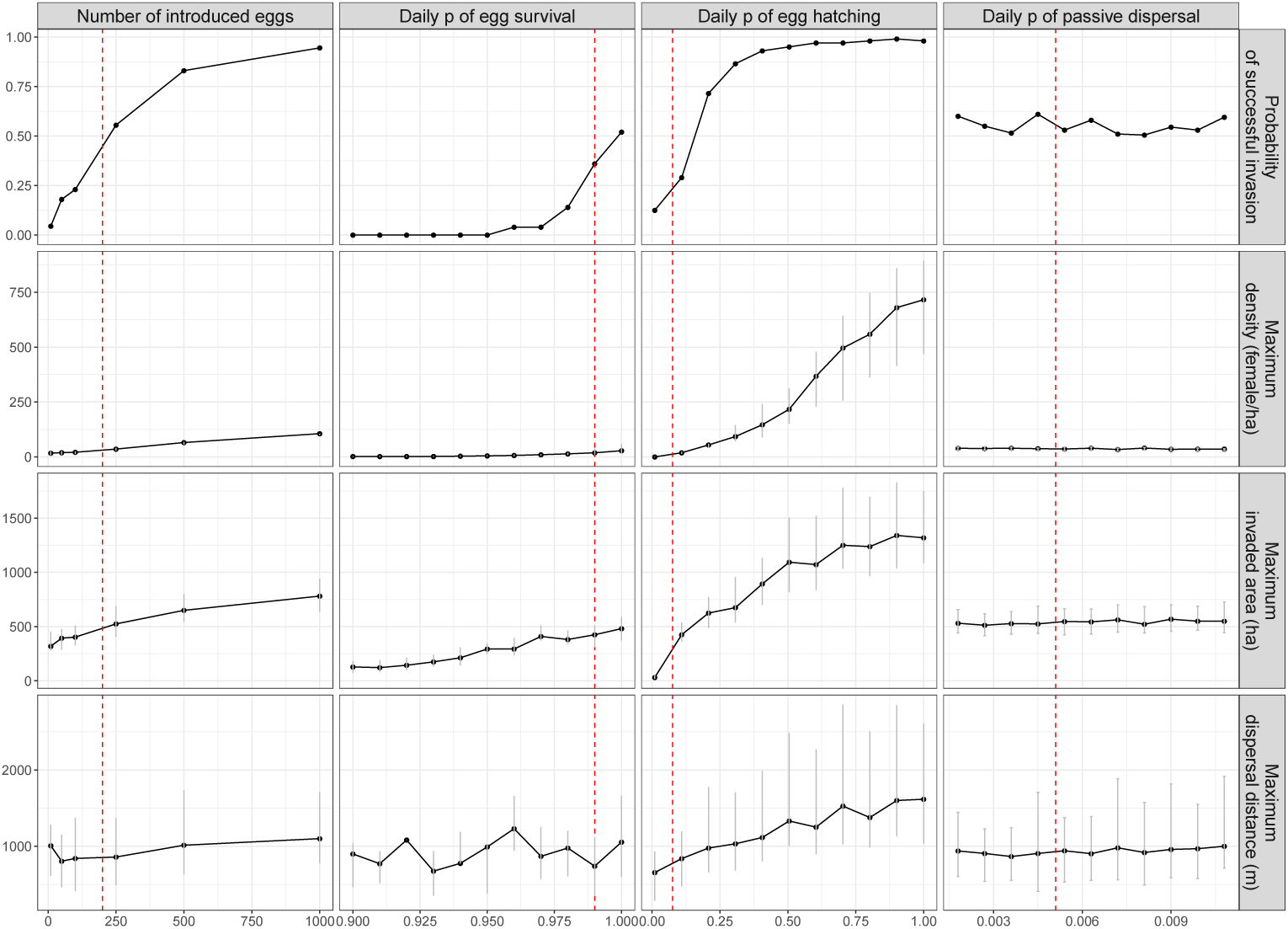
Matrix of scatterplots showing the effect of the four model parameters (column) selected for sensitivity analysis on different model outputs (rows). The black dots represent median values across the 200 iterations while vertical grey lines the inter-quartile range for each type of output, except for the probability of successful invasion which is cumulative across iterations. The red dashed vertical lines indicate the parameters value used in model simulations.

## References

Abozeid, S., Elsayed, A. K., Schaffner, F., and Samy, A. M. (2018). Re-emergence of *Aedes aegypti* in Egypt. The Lancet Infectious Diseases, 18(2):142–143.

Adhami, J. and Reiter, P. (1998). Introduction and establishment of *Aedes (Stegomyia) albopictus skuse* (Diptera: Culicidae) in Albania. Journal of the American Mosquito Control Association, 14(3):340–343.

Alemanno, A., Fiorello, D., Martino, A., Pasaoglu, G., Scarcella, G., Thiel, C., Zubaryeva, C., European Commission, Joint Research Centre, and Institute for Energy and Transport (2012). Driving and parking patterns of European car drivers a mobility survey. European Commission - Joint Research Centre, Institute for Institute for Energy and Transport. OCLC: 1111229299.

Beck, H. E., Zimmermann, N. E., McVicar, T. R., Vergopolan, N., Berg, A., and Wood, E. F. (2018). Present and future Köppen-Geiger climate classification maps at 1-km resolution. Scientific Data, 5:180214.

Blackburn, T. M., Cassey, P., and Duncan, R. P. (2020). Colonization pressure: a second null model for invasion biology. Biological Invasions, 22(4):1221–1233.

Brady, O. J. and Hay, S. I. (2019). The first local cases of Zika virus in Europe. The Lancet, 0(0).

Brown, J. E., Scholte, E.-J., Dik, M., Den Hartog, W., Beeuwkes, J., and Powell, J. R. (2011). *Aedes aegypti* mosquitoes imported into the netherlands, 2010. Emerging infectious diseases, 17(12):2335.

Caminade, C., Medlock, J. M., Ducheyne, E., McIntyre, K. M., Leach, S., Baylis, M., and Morse, A. P. (2012). Suitability of European climate for the Asian tiger mosquito *Aedes albopictus*: recent trends and future scenarios. Journal of The Royal Society Interface, 9(75):2708–2717.

Cardamatis, J. P. (1929). Dengue in Greece. Bulletin de la Société de Pathologie Exotique, 22(4):272–292.

Carrasquilla, M. C. and Lounibos, L. P. (2015). Satyrization without evidence of successful insemination from interspecific mating between invasive mosquitoes. Biological Letters, 11(9):20150527.

Center for International Earth Science Information Network - CIESIN - Columbia University and Information Technology Outreach Services - ITOS - University of Georgia (2013). Global Roads Open Access Data Set, Version 1 (gROADSv1).

Christophers, S. R. (1960a). Aedes aegypti (L.) The Yellow Fever Mosquito: Its Life History, Bionomics and Structure. Cambridge University Press.

Christophers, S. R. (1960b). XXII - Mating and oviposition, pages 510–511. Cambridge University Press.

Copernicus Climate Change Service (2017). Era5: Fifth generation of ecmwf atmospheric reanalyses of the global climate. copernicus climate change service climate data store (cds).

Copernicus project (2019). EU-DEM v1.1 — Copernicus Land Monitoring Service.

Dunn, A. M. and Hatcher, M. J. (2015). Parasites and biological invasions: parallels, interactions, and control. Trends in Parasitology, 31(5):189–199.

Díaz-Nieto, L. M., Chiappero, M. B., Astarloa, C. D. d., Maciá, A., Gardenal, C. N., and Berón, C. M. (2016). Genetic Evidence of Expansion by Passive Transport of *Aedes (Stegomyia) aegypti* in Eastern Argentina. PLOS Neglected Tropical Diseases, 10(9):e0004839. Publisher: Public Library of Science.

ECDC, E. C. f. D. P., Control, and Authority, E. F. S. (2019). The current (january 2019) known distribution of Aedes aegypti in europe at ‘regional’ administrative level. https://ecdc.europa.eu/en/disease-vectors/surveillance-and-disease-data/mosquito-maps.

Ekman, A. (2018). China in the mediterranean: An emerging presence. Institut Français Des Relations Internationales, pages 6–20.

Eritja, R., Escosa, R., Lucientes, J., Marques, E., Roiz, D., and Ruiz, S. (2005). Worldwide invasion of vector mosquitoes: present european distribution and challenges for spain. Biological invasions, 7(1):87.

Eritja, R., Palmer, J. R. B., Roiz, D., Sanpera-Calbet, I., and Bartumeus, F. (2017). Direct Evidence of Adult *Aedes albopictus* Dispersal by Car. Scientific Reports, 7(1):14399.

Eurostat (2018). Top 20 ports - volume (in TEUs) of containers handled in each port, by loading status (main ports).

Focks, D. A., Sackett, S. R., Bailey, D. L., and Dame, D. A. (1981). Observations on Container-Breeding Mosquitoes in New Orleans, Louisiana, with an Estimate of the Population Density of *Aedes aegypti* (L.). The American Journal of Tropical Medicine and Hygiene, 30(6):1329–1335.

Garcia, G. d. A., dos Santos, L. M. B., Villela, D. A. M., and Maciel-de Freitas, R. (2016). Using Wolbachia Releases to Estimate *Aedes aegypti* (Diptera: Culicidae) Population Size and Survival. PLoS One, 11(8).

Gloria-Soria, A., Lima, A., Lovin, D. D., Cunningham, J. M., Severson, D. W., and Powell, J. R. (2018). Origin of a High-Latitude Population of *Aedes aegypti* in Washington, DC. American Journal of Tropical Medicine and Hygiene, 98(2):445–452.

Ibañez-Justicia, A., Gloria-Soria, A., Den Hartog, W., Dik, M., Jacobs, F., and Stroo, A. (2017). The first detected airline introductions of yellow fever mosquitoes (*Aedes aegypti*) to europe, at schiphol international airport, the netherlands. Parasites & vectors, 10(1):603.

IMO, I. M. O. (2019). International shipping and world trade. facts and figures. http://www.imo.org/.

Kanamitsu, M., Ebisuzaki, W., Woollen, J., Yang, S.-K., Hnilo, J. J., Fiorino, M., and Potter, G. L. (2002). NCEP–DOE AMIP-II Reanalysis (R-2). Bulletin of the American Meteorological Society, 83(11):1631–1644.

Kearney, M. R., Gillingham, P. K., Bramer, I., Duffy, J. P., and Maclean, I. M. (2020). A method for computing hourly, historical, terrain-corrected microclimate anywhere on earth. Methods in Ecology and Evolution, 11(1):38–43.

Knudsen, A. B., Romi, R., and Majori, G. (1996). Occurrence and spread in Italy of Aedes albopictus, with implications for its introduction into other parts of Europe. Journal of the American Mosquito Control Association, 12(2):177–183.

Kraemer, M. U., Reiner, R. C., Brady, O. J., Messina, J. P., Gilbert, M., Pigott, D. M., Yi, D., Johnson, K., Earl, L., Marczak, L. B., et al. (2019). Past and future spread of the arbovirus vectors *Aedes aegypti* and *Aedes albopictus*. Nature microbiology, 4(5):854.

Kraemer, M. U. G., Sinka, M. E., Duda, K. A., Mylne, A. Q. N., Shearer, F. M., Barker, C. M., Moore, C. G., Carvalho, R. G., Coelho, G. E., Van Bortel, W., Hendrickx, G., Schaffner, F., Elyazar, I. R. F., Teng, H.-J., Brady, O. J., Messina, J. P., Pigott, D. M., Scott, T. W., Smith, D. L., Wint, G. R. W., Golding, N., and Hay, S. I. (2015). The global distribution of the arbovirus vectors *Aedes aegypti* and *Ae. albopictus*. Elife, 4:e08347.

Leta, S., Beyene, T. J., De Clercq, E. M., Amenu, K., Revie, C. W., and Kraemer, M. U. G. (2018). Global risk mapping for major diseases transmitted by *Aedes aegypti* and *Aedes albopictus*. International Journal of Infectious Diseases, 67:25–35.

Lindh, E., Argentini, C., Remoli, M. E., Fortuna, C., Faggioni, G., Benedetti, E., Amendola, A., Marsili, G., Lista, F., Rezza, G., et al. (2019). The italian 2017 outbreak chikungunya virus belongs to an emerging aedes albopictus–adapted virus cluster introduced from the indian subcontinent. In Open forum infectious diseases, volume 6, page ofy321. Oxford University Press US.

Liu, B., Gao, X., Ma, J., Jiao, Z., Xiao, J., Hayat, M. A., and Wang, H. (2019). Modeling the present and future distribution of arbovirus vectors *Aedes aegypti* and *Aedes albopictus* under climate change scenarios in mainland china. Science of The Total Environment, 664:203–214.

Liumbruno, G. M., Calteri, D., Petropulacos, K., Mattivi, A., Po, C., Macini, P., Tomasini, I., Zucchelli, P., Silvestri, A. R., Sambri, V., et al. (2008). The chikungunya epidemic in italy and its repercussion on the blood system. Blood transfusion, 6(4):199.

Lockwood, J. L., Cassey, P., and Blackburn, T. (2005). The role of propagule pressure in explaining species invasions. Trends in Ecology & Evolution, 20(5):223–228.

M. Friedl, D. S.-M. (2015). MCD12q1 MODIS/Terra+Aqua Land Cover Type Yearly L3 Global 500m SIN Grid V006. type: dataset.

Maclean, I. M. D., Mosedale, J. R., and Bennie, J. J. (2019). Microclima: An r package for modelling meso- and microclimate. Methods in Ecology and Evolution, 10(2):280–290.

Marcantonio, M., Metz, M., Baldacchino, F., Arnoldi, D., Montarsi, F., Capelli, G., Carlin, S., Neteler, M., and Rizzoli, A. (2016). First assessment of potential distribution and dispersal capacity of the emerging invasive mosquito *Aedes koreicus* in Northeast Italy. Parasites & Vectors, 9(1):63.

Marcantonio, M., Reyes, T., and Barker, C. M. (2019). Quantifying *Aedes aegypti* dispersal in space and time: a modeling approach. Ecosphere, 10(12):e02977.

Metzger, M. E., Hardstone Yoshimizu, M., Padgett, K. A., Hu, R., and Kramer, V. L. (2017). Detection and Establishment of *Aedes aegypti* and *Aedes albopictus* (Diptera: Culicidae) Mosquitoes in California, 2011-2015. Journal of Medical Entomology, 54(3):533–543.

Montecino, D., Marcantonio, M., Perkins, A., and Barker, C. (2016). Modeling *Aedes albopictus* Skuse population dynamics and spatial behavior in urban landscapes, an integrated approach. Parasites & Vectors.

Mordecai, E. A., Caldwell, J. M., Grossman, M. K., Lippi, C. A., Johnson, L. R., Neira, M., Rohr, J. R., Ryan, S. J., Savage, V., Shocket, M. S., et al. (2019). Thermal biology of mosquito-borne disease. Ecology letters, 22(10):1690–1708.

Nelson, A., Weiss, D. J., van Etten, J., Cattaneo, A., McMenomy, T. S., and Koo, J. (2019). A suite of global accessibility indicators. Scientific Data, 6(1):1–9.

Neteler, M., Bowman, M. H., Landa, M., and Metz, M. (2012). GRASS GIS: A multi-purpose open source GIS. Environmental Modelling & Software, 31:124–130.

Oliva, L. O., La Corte, R., Santana, M. O., and de Albuquerque, C. M. (2018). Quiescence in *Aedes aegypti*: Interpopulation differences contribute to population dynamics and vectorial capacity. Insects, 9(3):111.

Otero, M., Schweigmann, N., and Solari, H. G. (2008). A Stochastic Spatial Dynamical Model for *Aedes aegypti*. Bulletin of Mathematical Biology, 70(5):1297.

Pascoe, E. L., Pareeth, S., Rocchini, D., and Marcantonio, M. (2019). A Lack of “Environmental Earth Data” at the Microhabitat Scale Impacts Efforts to Control Invasive Arthropods That Vector Pathogens. Data, 4(4):133. Number: 4 Publisher: Multidisciplinary Digital Publishing Institute.

Powell, J. R., Tabachnick, W. J., Powell, J. R., and Tabachnick, W. J. (2013). History of domestication and spread of *Aedes aegypti* - A Review. Memórias do Instituto Oswaldo Cruz, 108:11–17.

R Core Team (2019). R: A Language and Environment for Statistical Computing. R Foundation for Statistical Computing, Vienna, Austria.

Reinhold, J. M., Lazzari, C. R., and Lahondère, C. (2018). Effects of the environmental temperature on *Aedes aegypti* and *Aedes albopictus* mosquitoes: a review. Insects, 9(4):158.

Rezza, G., Nicoletti, L., Angelini, R., Romi, R., Finarelli, A. C., Panning, M., Cordioli, P., Fortuna, C., Boros, S., Magurano, F., Silvi, G., Angelini, P., Dottori, M., Ciufolini, M. G., Majori, G. C., Cassone, A., and CHIKV study group (2007). Infection with chikungunya virus in Italy: an outbreak in a temperate region. Lancet, 370(9602):1840–1846.

Richards, S. L., Ponnusamy, L., Unnasch, T. R., Hassan, H. K., and Apperson, C. S. (2006). Host-feeding patterns of *Aedes albopictus* (Diptera: Culicidae) in relation to availability of human and domestic animals in suburban landscapes of central North Carolina. Journal of Medical Entomology, 43(3):543–551.

Ritchie, S. A., Montgomery, B. L., and Hoffmann, A. A. (2013). Novel Estimates of *Aedes aegypti* (Diptera: Culicidae) Population Size and Adult Survival Based on Wolbachia Releases. Journal of Medical Entomology, 50(3):624–631.

Roche, B., Léger, L., L’Ambert, G., Lacour, G., Foussadier, R., Besnard, G., Barré-Cardi, H., Simard, F., and Fontenille, D. (2015). The Spread of *Aedes albopictus* in Metropolitan France: Contribution of Environmental Drivers and Human Activities and Predictions for a Near Future. PLoS ONE, 10(5):e0125600.

Ryan, S. J., Carlson, C. J., Mordecai, E. A., and Johnson, L. R. (2019). Global expansion and redistribution of aedes-borne virus transmission risk with climate change. PLoS neglected tropical diseases, 13(3).

Sabatini, A., Raineri, V., Trovato, G., and Coluzzi, M. (1990). *Aedes albopictus* in Italy and possible diffusion of the species into the Mediterranean area. Parassitologia, 32(3):301–304.

Schaffner, F. and Mathis, A. (2014). Dengue and dengue vectors in the WHO European region: past, present, and scenarios for the future. The Lancet Infectious Diseases, 14(12):1271–1280.

Schneider, A., Friedl, M. A., and Potere, D. (2009). A new map of global urban extent from MODIS satellite data. Environmental Research Letters, 4(4):044003.

Service, M. W. (1997). Mosquito (Diptera: Culicidae) dispersal–the long and short of it. Journal of Medical Entomology, 34(6):579–588.

Simberloff, D. (2014). Biological invasions: What’s worth fighting and what can be won? Ecological Engineering, 65:112–121.

Smith, A., Lott, N., and Vose, R. (2011). The Integrated Surface Database: Recent Developments and Partnerships. Bulletin of the American Meteorological Society, 92(6):704–708.

Soares-Pinheiro, V. C., Dasso-Pinheiro, W., Trindade-Bezerra, J. M., and Tadei, W. P. (2016). Eggs viability of *Aedes aegypti* Linnaeus (Diptera, Culicidae) under different environmental and storage conditions in Manaus, Amazonas, Brazil. Brazilian Journal of Biology, 77(2):396–401.

Tsunoda, T., Cuong, T. C., Dong, T. D., Yen, N. T., Le, N. H., Phong, T. V., and Minakawa, N. (2014). Winter refuge for *Aedes aegypti* and *Ae. albopictus* mosquitoes in hanoi during winter. PloS one, 9(4).

Vanwambeke, S. O., Linard, C., and Gilbert, M. (2019). Emerging challenges of infectious diseases as a feature of land systems. Current Opinion in Environmental Sustainability, 38:31–36.

Venturi, G., Luca, M. D., Fortuna, C., Remoli, M. E., Riccardo, F., Severini, F., Toma, L., Manso, M. D., Benedetti, E., Caporali, M. G., Amendola, A., Fiorentini, C., Liberato, C. D., Giammattei, R., Romi, R., Pezzotti, P., Rezza, G., and Rizzo, C. (2017). Detection of a chikungunya outbreak in Central Italy, August to September 2017. Eurosurveillance, 22(39):17–00646.

Yang, H., Macoris, M. d. L. d. G., Galvani, K., Andrighetti, M., and Wanderley, D. (2009). Assessing the effects of temperature on the population of *Aedes aegypti*, the vector of dengue. Epidemiology & Infection, 137(8):1188–1202.

